# scEvoNet: a gradient boosting-based method for prediction of cell state evolution

**DOI:** 10.1101/2022.12.07.519467

**Authors:** Aleksandr Kotov, Andrei Zinovyev, Anne-Helene Monsoro-Burq

## Abstract

**Background:** Exploring the function or the developmental history of cells in various organisms provides insights into a given cell type’s core molecular characteristics and putative evolutionary mechanisms. Numerous computational methods now exist for analyzing single-cell data and identifying cell states. These methods mostly rely on the expression of genes considered as markers for a given cell state. Yet, there is a lack of scRNA-seq computational tools to study the evolution of cell states, particularly how cell states change their molecular profiles. This can include novel gene activation or the novel deployment of programs already existing in other cell types, known as co-option.

**Results:** Here we present scEvoNet, a Python tool for predicting cell type evolution in crossspecies or cancer-related scRNA-seq datasets. ScEvoNet builds the confusion matrix of cell states and a bipartite network connecting genes and cell states. It allows a user to obtain a set of genes shared by the characteristic signature of two cell states even between distantly-related datasets. These genes can be used as indicators of either evolutionary divergence or co-option occurring during organism or tumor evolution. Our results on cancer and developmental datasets indicate that scEvoNet is a helpful tool for the initial screening of such genes as well as for measuring cell state similarities.

**Conclusions:** The scEvoNet package is implemented in Python and is freely available from https://github.com/monsoro/scEvoNet. Utilizing this framework and exploring the continuum of transcriptome states between developmental stages and species will help explain cell state dynamics.

## Background

Cells, the fundamental construction blocks of multicellular organisms, are characterized by great diversity in complex multicellular organisms. They include differentiated and function-specific cells, their stem cells for cell renewal during lifetime, and all the transitional states between these two points. In disease, cell and tissue homeostasis are altered, leading to the appearance of new pathological and dysfunctional cells. During evolution, the diversification of cell types is caused by genomic individualization relying on fundamental evolutionary principles such as functional segregation, divergence, co-option of gene modules, and *de novo* gene emergence. Co-option of gene programs is a mechanism allowing the emergence of new functions in a cell type by using existing gene networks from other cell types [1, 2]. The understanding of cell biology emanates from describing cells by their functions, their gene expression, interactions with their environment, and their lineage relationships. The emergence of single-cell RNA sequencing (scRNA-seq) began a new age of transcriptomic research, extending our understanding of cell heterogeneity and dynamics. Highly detailed atlases of cell types were produced for many tissues and organisms, in normal or pathological conditions [3–7]. Comparing those highly divergent datasets would allow asking key questions regarding the conservation of core genetic programs in poorly-related cellular contexts, the origins of cellular diversity and its evolutionary mechanisms, or the transcriptional paths leading to disease. However, data received from various biological conditions and various organisms is entangled by technical and biological batch effects which vastly complicates their comparison [8, 9]. Thus, forces shaping transcriptome dynamics remain poorly understood. Another application of scRNA-seq in evolutionary biology is accessing tumor heterogeneity and tracking its transformation as well as assessing the selective evolution of tumors during therapy or metastatic progression [10]. ScRNA-seq overcomes the constraints of classic bulk RNA sequencing by estimating transcriptome at a single-cell level and characterizing various cell types in the tumor microenvironment. Moreover, this allows a better understanding of the molecular mechanisms facilitating tumor occurrence. Although it could potentially reveal the somatic mutations during tumor evolution, scRNA-seq data sparsity [11] often prevents mutation calling (one of the main information sources for studying tumor evolution). Still, the scRNA-seq of tumors can determine the dynamic changes in tumor heterogeneity and the transcriptional evolution of tumor cells during metastasis development [12].

Currently, there is a lack of a specific tool that uses closely or distantly-related scRNA-seq datasets as input to study the potential co-option and evolution of gene programs between different organisms during development and differentiation, or between tumor cells at different stages of tumor progression. Kun Xu et al. [12] used Monocle [13] and scVelo [14] to study transcriptome dynamics of malignant cells between the primary tumor and lymph node metastases. They also used NATMI [15] for the generation of the cell-to-cell receptor-ligand network where edges are generated based on the expression of the ligand in one cell type and of its related receptor in another cell type. However, this latter strategy is not designed to study co-option in cancer which is a crucial mechanism forcing the molecular changes that propel tumor progression [16]. In another work, Pandey et al. use scRNA-seq to study the evolutions of neuronal types by comparing cell types in larva and adult zebrafish. They utilized Random Forest to generate a model for each cell type and predict cells with each model to build a confusion matrix mapping cell types by the number of cells predicted with each model [17]. Yet, this strategy is not wrapped into the usable framework and cannot be used to extract genes that are characteristic of cell type transitions. We present scEvoNet, a method that builds a cell type-to-gene network using the Light Gradient Boosting Machine (LGBM) algorithm [18] overcoming different domain effects (different species/different datasets) and dropouts that are inherent for the scRNA-seq data [19]. This tool predicts potentially co-opted genes together with genes characteristic of each cell state during development across species. Recently we showed the ability of a similar LGBM-based classifier to detect neural crest cells in distantly-related scRNA-seq datasets [20]. Despite technical batch effects (datasets were made in different laboratories with different technologies) and biological batch effects (datasets were from two evolutionarily distant organisms and at different developmental time points), we have achieved a high AUC score of 0.95 for classifying zebrafish cells with our frog-based NC model [20]. Here we have expanded this method: scEvoNet applies to a variety of applications, e.g., between different time points during a given organism’s development, between species, and when comparing primary tumor and metastasis. We believe that scEvoNet will facilitate the study of cell state transitions in a variety of contexts and from highly divergent datasets.

## Implementation

The workflow of scEvoNet is illustrated in Fig. 1A. scEvoNet takes (1) an expression matrix, and (2) a list of cell labels as input data per organism/time point of interest. For each cell type provided by a user, scEvoNet generates an LGBM binary classifier (one cell type vs all other cells) in two steps (Fig. 1B). Firstly, it generates a model considering all genes in the dataset. For the obtained model of the particular cell type, scEvoNet selects the top 3000 important features (cell types related genes) and uses only them to re-train the final cell type model which will be used for the generation of the cell types confusion matrix. This is part of the domain adaptation that we perform to make the resulting model less dependent on the number of genes that are missing in another dataset to which the model will be applied. Additionally, to reduce the effect of the different biological domains between datasets and to reduce the effect of the scRNA-seq data sparsity we apply a sigmoid function that smooths expression units more flexible than simple binarization, which has been shown to keep enough information for scRNA-seq data analysis [21]. For the model training, we use early stopping to avoid overfitting with 10 rounds which determines the actual number of estimators in the regressions.

**Figure 1.**
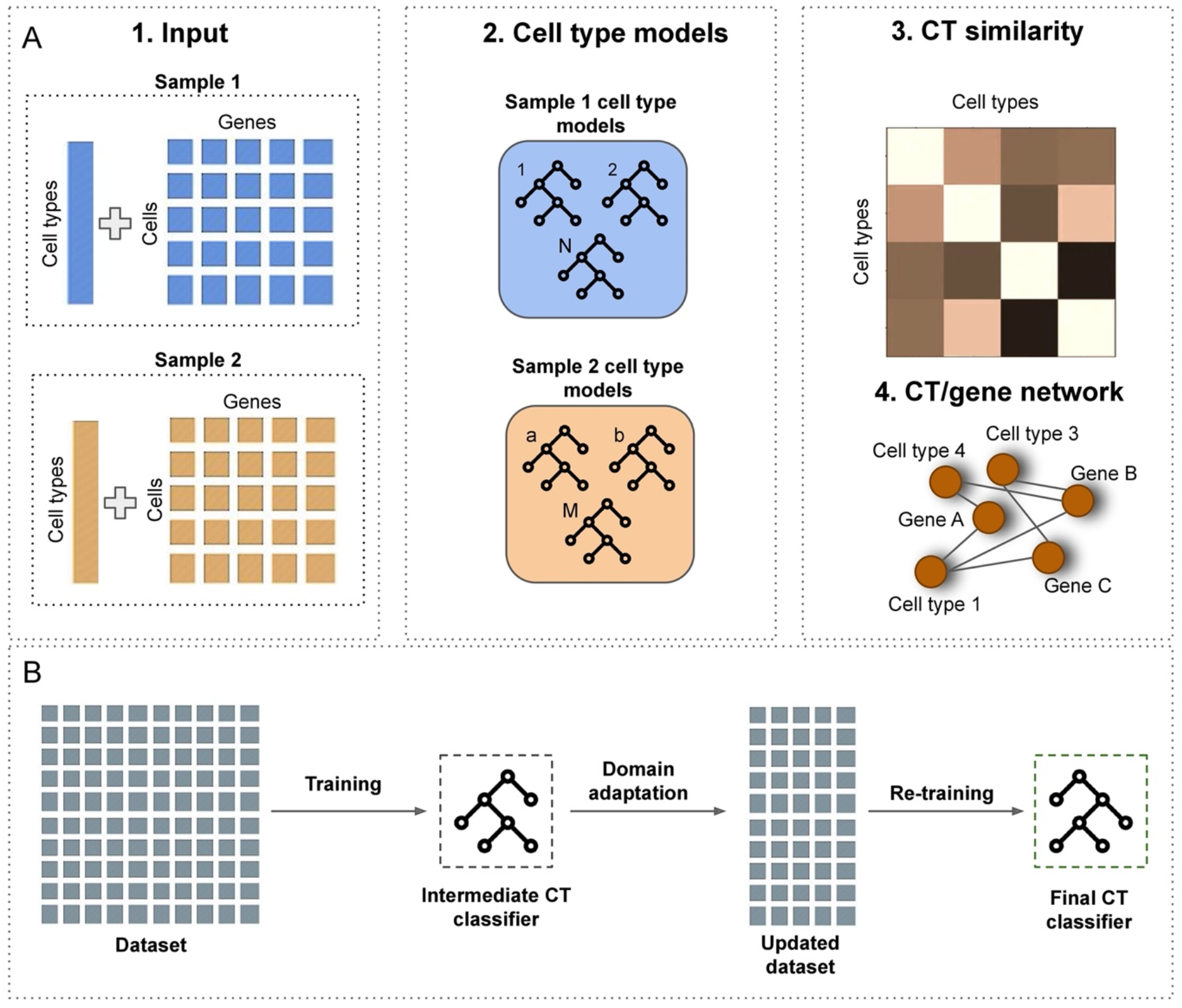
scEvoNet scheme. A) scEvoNet takes a list of clusters and a matrix of expressions for each sample as input. For each sample, it generates an object with cell type classifiers and top important features for each cluster from the provided set of clusters. In the final step, the tool builds a confusion matrix and a network of genes associated with each cell type. B) We use the LGBM algorithm to produce a classifier for each cell type. To smooth the data in order to deal with the batch effects we apply the sigmoid function and only the top important features to create the final model.

Next, scEvoNet uses each cell type model to predict cells from both datasets. This way we get a confusion matrix with cell type to cell type comparison. In the next step, scEvoNet builds a network where the nodes are cell types or genes so that cell types can only connect to genes. To do this, firstly we extract top features (genes) which are important for each cell type (both positively correlated and negatively correlated) and then combine all the cell type-related top important genes into one main network with all cell types and cell type-related genes. This strategy is similar to GRNboost2 [22] which outperformed many other tools in a recent benchmarking study [23]. GRNboost2 generates a gene-gene network similarly, whereas scEvoNet extends it to all cell types in two datasets. Furthermore, scEvoNet implements a shortest path search in order to generate a subnetwork of interest. For example, to study the evolution of a particular cell type, a user might request all the shortest paths (with a selected cut-off on their length) between two cell types and scEvoNet will yield all the genes and cell types between these two using the confusion matrix as a metric of the cell types similarity. Each gene-to-cell type connection has an importance value (a score displaying how useful each feature was in the building of the boosted decision trees within the model) by which users can filter sub-networks.

## Results

First, we applied scEvoNet to identify core characteristics during the evolution of the neural crest (NC) cells using two different vertebrate organisms. The NC is a multipotent and migratory cell population unique to vertebrates and essential notably for pigment, peripheral and enteric nervous system, and craniofacial structures formation [24]. For the input data, we used whole embryo scRNA-seq datasets for the *Xenopus tropicalis*, a non-amniote tetrapod vertebrate, at an early developmental stage (late gastrulation stage 12, and neurulation stages 13 (neural plate) and 14 (neural fold)) and *Mus musculus*, a mammal, at a similar developmental stage (late gastrulation stage 8.25) [4, 25].

Before using scEvoNet, we validated how one NC-based classifier trained on the *Xenopus* dataset will recognize NC cells in the whole embryo mouse dataset and obtained the result of a 0.89 AUC score (Fig. 2A, B). Next, using published cluster annotations for these two whole embryo scRNA-seq datasets, we run scEvoNet and obtained the confusion matrix (Fig. 2C). In this dataset of extended complexity, the highest similarity score for frog NC remains the mouse NC. Thereafter, we built a network of cell types and related genes. To identify genes that are highly conserved and cell type-specific in the evolution of the NC we selected a sub-network that consists of the shortest paths from *Xenopus* NC to mouse NC with the top 3 close cell types according to the confusion matrix. To obtain a larger subnetwork number_of_shortest_paths=300 was used. Subsequently, we determined several groups of genes differently related to cell types within this subnetwork (Fig. 2D). The first group includes genes that are associated between NC and a closely-related cell type in only one organism (e.g. NC and neural plate in frog: *sox2, sox3, snai2, hes1, zic1;* NC and midhindbrain in mice: *gadd45a, mdk, ptn*). If this gene expression signature was the consequence of the evolutionary divergence of function, this could be studied using the scRNA-seq of the ancestor organisms. The second group consists of genes that are characteristic of NC both in *Xenopus* and *Mus musculus (pax3, tfap2c, tfap2a, tfap2b, sox9):* all are known markers of NC or their progenitors [26]. The third group includes genes that are associated between the frog NC and mouse NC, and shared with the mouse NC-related cell types (as defined with confusion matrix of similarities): *mafb, cldn6* for the neural plate; *zic3, tfap2a* for the midhindbrain. Thus, our tool was not only able to construct a matrix of similar cell types that can be used to study cell types similarities, but also defined three groups of genes that may have diverse roles in a crossspecies transformation of the molecular profile of neural crest cells.

**Figure 2.**
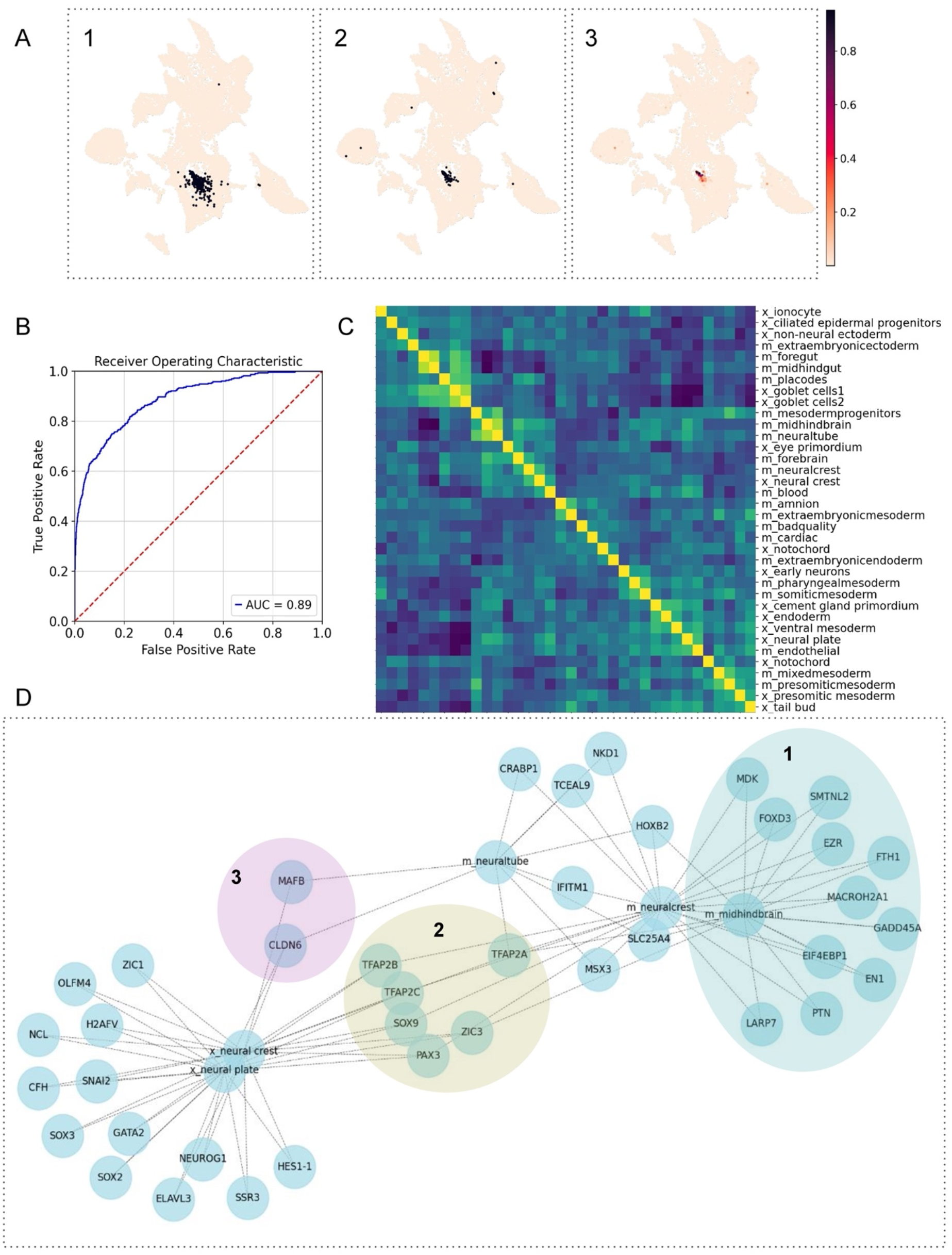
The development of the specific cell type between frog and mouse. A) First UMAP represents highlighted annotated neural crest cells in the whole embryo dataset, second UMAP represents predicted neural crest cells with our classifier, and third UMAP represents predicted scores of our classifier. B) The AUC score for the neural crest classifier is 0.89 C) The confusion matrix for mouse and frog samples (prefix x_ is for *Xenopus*, prefix m_ is for mouse). The values in the confusion matrix are the correlations between two lists of scores for all cell type models. D) Selecting the subnetwork of 300 shortest paths from *Xenopus* neural crest to mouse neural crest shows genes that are shared with closely-related cell types, such as *mafb* or *cldn6* (group 3). It reveals two groups of genes: genes from group 1 are organism-specific genes, and genes from group 2 are important genes for the specific cell type (NC) in both organisms.

Next, we applied scEvoNet to a human breast cancer metastasis dataset [12]. We selected a patient with available datasets for the primary tumor and the lymph node metastases. We used a standard Scanpy [27] pipeline to obtain clusters for both matrices (primary tumor and metastasis). Marker genes from the source paper were used to annotate obtained clusters (Fig. 3A). First, using scEvoNet, we calculated the confusion matrix (Fig. 3B). As expected, we observed high connectivity between cells of the same type from the primary tumor or metastasis in lymph nodes, e.g., B cells, plasma cells, immune cells, macrophages, dendritic cells, and tumor cells. Next, to study cancer cell evolution, we used scEvoNet to discover common cluster-specific genes between the most distant malignant clusters from the primary tumor (cluster p_cancer_cells_cox6+) and metastasis (cluster m_cancer_cells_gapdh+). We found 14 genes that were directly connected to both cell types (Fig. 3C), among them *malat1*, levels of which inversely correlate with breast cancer progression and metastatic capacity [28], and *b2m* an important marker involved in carcinogenesis, invasion, and metastasis [29]. Among this list of genes directly connecting two tumor cell types were several mitochondrial genes. Although a common hypothesis relates the expression of mitochondrial genes to sample or data processing artifacts, growing evidence supports the importance of mitochondrial genes in cancer metastasis [30]. Next, we explored what other genes from close cell types might be involved in tumor evolution. To do so, we selected a subnetwork with all the genes related to cell types of interest and 5 similar cell types according to the confusion matrix obtained earlier. As a result, we determined two cancer cell types (metastatic and primary) that have as a common network neighbor the lymph node B cells, through genes *hmgb1* and *b2m*. Interestingly, it was shown previously that exosomal *hmgb1* promotes hepatocellular carcinoma immune evasion by stimulating TIM-1+ regulatory B cell expansion [31]. Also, blockade of the *hmgb1* signaling pathway inhibits tumor growth in diffuse large B-cell lymphoma [32]. In another work, *b2m* specific B cells were defined as the most important prometastatic B cell cluster essentially contributing to distant metastasis in Clear Cell Renal Cell Carcinoma [33]. *B2m* is also an important element in the immune escape mechanism since a decrease in *b2m* expression reduces the number of antigens presented on the cell surface, including tumor-related antigens, which has been shown in particular in diffuse large B-cell lymphoma [34]. Thus, scEvoNet here provides a result supported by the literature, suggesting that users can retrieve meaningful gene candidates involved in tumor progression and immune escape in cancer.

**Figure 3.**
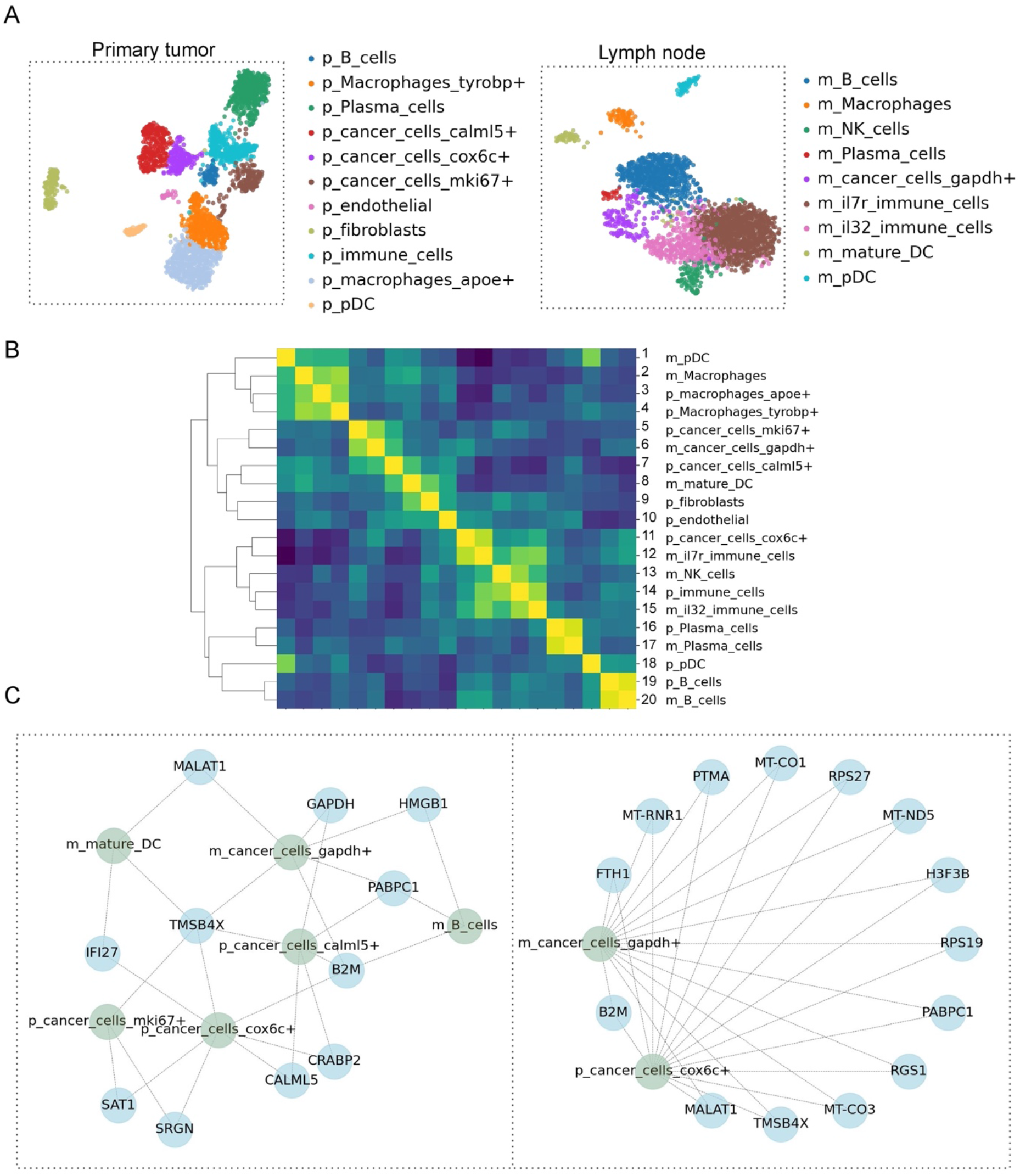
Primary tumor vs metastasis comparison. A) UMAPs for the primary human breast cancer (left) and metastasis in the lymph node (right). B) The confusion matrix shows different rates of similarity between different clusters of cancer cells in primary tumor and metastasis (the p_ prefix is for primary, and the m_ prefix is for metastasis). The values in the confusion matrix are the correlations between two lists of scores for all cell type models. C) Two subnetworks of the relation of the cluster of cancer cells in primary tumor and cancer cells in metastasis. On the left subnetwork, we show only genes related to some other cell types, on the right subnetwork we selected genes that are directly connected to clusters of interest.

## Conclusions

The evolution of cell types and gene programs is one of the main focuses of developmental biology and is crucial for a better understanding of the origin of particular functions. For the moment, there is a lack of computational tools to address this question using the abundant scRNA-seq data publicly available databases. One existing approach is not wrapped into the usable framework (e.g., R/Python package) and has only one application (cell states comparison) so it cannot be used to extract genes that are responsible for cell types transitions such as co-opted genes or genes conservatively important for several cell states [17]. In this manuscript, we present scEvoNet, a method for analyzing the evolution of cell states from highly sparse scRNA-seq data. We show that it is applicable to studying cross-species and tumor-to-metastasis transitions. With this tool, we re-discover a canonical gene signature that remains conserved through evolution, and also predict species-specific genes and new candidates associated with similar cell types. Our findings may indicate the co-option of genes or shared programs in closely related cell types. It also suggests the potential use of an immune escape mechanism in breast cancer metastasis, which has previously been shown in another cancer type. Yet, one limitation is that scEvoNet does not match gene sequences and only works with labels provided by the user, which can reduce the number of genes to be found between different cell types in cross-species comparison.

The tool is adjustable and can be utilized for an initial screening strategy. It is compatible with AnnData object format used in the Scanpy Python package [27].

## Availability and requirements

List the following:

Project name: scEvoNet
Project home: https://github.com/monsoro/scEvoNet
Operating system(s): Mac OS, Linux, Windows
Programming language: Python
License: MIT license
Any restrictions to use by non-academics: MIT license

## List of abbreviations

scRNA-seq: single-cell RNA sequencing
LGBM: Light Gradient Boosting Machine
NC: neural crest

## Declarations

### Ethics approval and consent to participate

Not applicable.

### Consent for publication

Not applicable.

### Availability of data and materials

We have been using public data and did not produce sequence data by ourselves.

### Competing interests

AZ is employed by EvoTec company.

### Funding

This project received funding from European Union’s Horizon 2020 research and innovation program under Marie Skłodowska-Curie grant agreement No 860635, NEUcrest ITN (to AHMB); Agence Nationale pour la Recherche (ANR-21-CE13-0028; to AHMB) and Institut Universitaire de France (to AHMB); Agence Nationale de la Recherche as part of the “Investissements d’avenir” program, reference ANR-19-P3IA-0001 (PRAIRIE 3IA Institute).

### Authors’ contributions

AK, AHMB, AZ conceived the project. AK designed the strategy, performed the work, did the programming, wrote and edited the manuscript. AZ reviewed the code and edited the manuscript. AHMB supervised the project and edited the manuscript.

## Acknowledgments

The authors are grateful to Drs. Igor Adameyko and Leon Peshkin for insightful scientific discussions and comments on the manuscript.

## Authors’ information

-

## Footnotes

-

## Notes

### Competing Interest Statement

AZ is an employee of Evotec

